# Pathogenicity Validation of *Parastagonospora pseudonodorum* on Wheat and Barley

**DOI:** 10.64898/2026.10.17.759865

**Authors:** Dianevys González-Peña Fundora, Mohamed Hafez, Mouldi Zid, Sean Wakowiak, Reem Aboukhaddour

## Abstract

*Parastagonospora pseudonodorum*, previously known as *Phaeosphaeria avenaria* f. sp. *tritici* (Pat1), is a fungal species within the Septoria blotch complex that causes yield losses in cereal crops worldwide. First reported in Canada in 1942 under an earlier taxonomic name. However, successive taxonomic changes in addition to its close resemblance to *Parastagonospora nodorum*, the causal agent of Septoria nodorum blotch have complicated its identification and its role in pathogenicity. Recent molecular identification of *P. pseudonodorum* in Western Canadian wheat samples necessitated the re-evaluation of its pathogenicity and raised concerns about potential misidentification with the closely-related species *P. nodorum.* These two species secrete combinations of various necrotic effectors (NEs) that contribute to their virulence. In this study, we examined the pathogenicity of *P. pseudonodorum* following Koch’s postulates. Our results confirmed the ability of *P. pseudonodorum* to infect the leaves and spikes of both hosts, but the symptoms was weaker on the leaves of both hosts and wheat spikes, whereas more evident damage was observed on barley spikes. While a limited number of genotypes was tested as a proof of concept, but the observed differences in symptom severity suggest host and tissue associated variation in susceptibility. These results highlight the need to expand our monitoring of the leaf spot complex to include this species and to further explore its host range and genomic characterization. Awareness of this pathogen is essential, particularly given its association with cereal spikes and its quarantine status in some countries which could impacts the export of Canadian grains overseas.

## Introduction

The genus *Parastagonospora* constitutes an important group of fungi in the family Phaeosphaeriaceae (order Pleosporales, class Dothidomycetes), with 31 species currently listed in Index Fungorum (2026). These species are distributed worldwide as pathogens or saprobes on crops and grasses, and several are major pathogen of cereal crops, including wheat, barley, oat, and maize (Gao et al. 2024). Historically, cereal pathogens in this group were classified under the genus *Septoria* (Cunfer, 2000). However, their taxonomy has undergone considerable revision, resulting in the reassignment of several species to the genus *Stagonospora* (teleomorph: *Phaeosphaeria*, syn. *Leptosphaeria*). This separation was based on conidial morphology: species in the genus *Septoria* have conidia more than 10 times as long as wide, whereas those in *Stagonospora* has conidia less than 10 times as long as they were wide (Sprague 1950). In 2013, the genus *Stagonospora* and its teleomorph *Phaeosphaeria* were renamed *Parastagonospora* to comply with the new rules of the International Code of Nomenclature for algae, fungi, and plants, which require a single name for pleomorphic fungi (those with both sexual and asexual forms) (Quaedvlieg et al. 2013).

The genus *Parastagonospora* includes several cereal pathogenic species and groups that were originally distinguished based on their conidia morphology, host specialization and formation of sexual structure (Shaw 1957a, 1957b). *Parastagonospora nodorum* is the most extensively studied species in the genus. It is a heterothallic fungus and a major pathogen causing Septoria nodorum blotch (SNB) of cereals, with wheat being the most susceptible host. *Parastagonospora avenae* was identified as a pathogen of oat and follows a heterothallic mating type behaviour. Within *P. avenae*, however, a group of isolates was identified by its inability to infect oat, but were weak pathogens of wheat; these isolates were homothallic and subsequently were designated as a distinct group, *Parastagonospora avenaria* f. sp. *triticea* (Pat) (Johnson 1947; Ueng et al. 2003). Pat isolates were first described from wheat and other cereals in Canada (Johnson 1947; Shaw 1957a, 1957b). Subsequent studies subdivided Pat into three groups (namely: Pat1, Pat2 and Pat3) based on variation in the internal transcribed spacer (ITS) sequence (Ueng et al. 2003; Ueng et al. 1998). Pat1 has been the most recovered group from wheat, while Pat2 was associated with foxtail barley, and Pat3 was also collected from wheat (Ueng and Chen, 1994; Ueng et al. 1995; Ueng et al. 2003; Ueng et al. 1998). More recently, Pat1was given a new designation as *Parastagonospora pseudonodorum* (Croll et al. 2021).This is a homothallic species has been reported primarily from wheat grains and spikes worldwide, including North America, and is considered a sister species to *P. nodorum* (Hafez et al. 2020; McDonald et al. 2012; Ueng et al. 2003).

Building on earlier evidence for a species complex of *Parastagonospora* species with cereals and grasses, McDonald et al. (2012) used phylogenetic analyses to show that this complex contains multiple cryptic species. Within the Pat complex, a total of six groups, Pat1-Pat6, were recognized expanding on the three previously described groups. Pat4, Pat5, and Pat6 were mainly isolated from wild grasses in Iran (Ghaderi et al. 2022; McDonald et al. 2012). Among the Pat isolates recovered from wheat, Pat1 had the most prevalent group and was reported on wheat grains and spikes worldwide, including North America (Hafez et al. 2020; McDonald et al. 2012). Two additional wheat-associated, but more distant lineages, P1 and P2, were isolated from wheat ears in Golestan Province in Iran and were clearly separated from the other *Parastagonospora* spp. across all loci examined and were interpreted as “undescribed relatives” within the Phaeosphaeriaceae rather than members of the Pat complex (McDonald et al. 2012; Zhang et al. 2009). Collectively, these findings illustrate that the traditional Pat designation encompasses several distinct evolutionary lineages within a broader *Parastagonospora* species complex consisting of *P. nodorum* and its relatives (McDonald et al. 2012).

Despite the taxonomic recognition of *P. pseudonodorum* as a distinct species in the cereal leaf spot complex, its pathogenicity and host/tissue specificity remain poorly resolved. Historical reports have differed in their assessment of its importance as pathogen of wheat and barley, its occurrence in leaves versus grains and spikes, and its geographical distribution (Johnson 1947; King et al. 1983; Richardson and Noble, 1970; Shaw 1957a; Shipton et al. 1971). Some of these discrepancies may reflect difficulties of distinguishing *P. pseudonodorum* from the closely related *P. nodorum*, a challenge that may persist even with molecular approaches when the markers used lack sufficient discriminatory power. Consequently, the pathogenicity and tissue specificity of *P. pseudonodorum* remain unresolved. In this study, we used morphologically and molecularly validated and identified isolates of *P. nodorum* and *P. pseudonodorum* to revisit Koch’s postulates and evaluate the ability of *P. pseudonodorum* to infect wheat and barley tissues, including spikes and leaves.

## Materials and methods

### Fungal culture

In this study, we used a single-spore isolates previously recovered in our Lab from wheat samples collected in Alberta (Hafez et al. 2020). The *P. pseudonodorum* isolates, C106-15 and B315-11, were recovered from the grains, while the *P. nodorum* isolate, C209-22a, was obtained from the nodes. These isolates secrete a combination of the following effectors: ToxA, Tox1, and Tox3. While C106-15 produces none of these effectors, B315-11 secretes Tox1, and C209-22a produces all three toxins. Isolate C106-15 was sequenced using Illumina short-read technology and identified as *P. pseudonodorum* by a species-specific qPCR assay and phylogenetic analysis ((Hafez et al. 2026) accession no. SAMN48453797). This identity was validated further by comparative whole-genome analysis against 2,668 Dothideomycete genomes from NCBI using NUCmer (Walkowiak et al. unpublished). Specifically, C106-15 shared 97.9 to 100% average nucleotide identity with reference *P. pseudonodorum* isolates, compared to only 69.6 to 69.9% with *P. nodorum* isolates, conclusively confirming its assignment as *P. pseudonodorum* rather than *P. nodorum* (Walkowiak et al. unpublished).

### Plant Material and Inoculation

Six wheat cultivars (Glenlea, Salamouni, Sumai3, Chinese Spring, Brandon, and Fielder) and 42 barley genotypes were sown in 10-cm pots containing Sunshine potting mix (W.R. Grace and Co.) at four seeds per pot. Plants were maintained in a growth cabinet at 20/18°C (day/night) with a 16-h photoperiod. All treatments were performed at least in triplicate. Inoculations were performed at the two-leaf stage (12 days post-sowing) to evaluate foliar infection. Additionally, a subset of four wheat and two barley genotypes (Glenlea, Salamouni, Chinese Spring, Fielder, Steptoe, and Rivers) was inoculated at anthesis (approx. 6-10 weeks post-sowing) to assess *P. pseudonodorum* ability to infect the spikes.

Spore suspension of each isolate was prepared following Liu et al (2004) by growing the colonies on 9-cm Petri dishes containing 25 mL of V8-PDA media (10 g/L of PDA, 15% of V8 juice, 3 g/L of CaCO_3_ and10 g/L of bacteriological agar), at room temperature (∼22°C) for seven days under continuous light to induce sporulation. The growing cultures were then flooded with autoclaved distilled water (dH_2_O) and the spores were scraped with a glass rod, and 20 uL of the fungal suspension were transferred to new V8-PDA plates and kept up to five days under continuous light to increase spore production. The spores were then filtrated through a Miracloth filter to eliminate mycelium debris, and the concentration was estimated using a Neubauer counting chamber. Plants were sprayed until run-off with a spore suspension adjusted to 10^6^ spores/mL and one drop of Tween 20 per 100 mL. Immediately following inoculation, the seedlings or spikes were covered with humid plastic bags and left at room temperature for 48 h before placing them back in the growth cabinet at 20/18°C (day/night) with a 16-h photoperiod. Disease severity on the second leaf was assessed 10 days post-inoculation (dpi) using a 1 to 5 scale based on Lamari & Bernier (1989). Symptoms on the spike were evaluated 18 dpi based on the number of infected spikelets per spike, and the intensity of infection based on the lesion colors. The bioassays were repeated two independent times.

### Fulfillment of Koch’s Postulates

All steps of Koch’s postulates, from inoculation and symptom association, isolation, re-inoculation, re-isolation and confirmation of symptoms were carried out to assess *P. pseudonodorum* ability to infect wheat/barley at 2ed leaf stage and on the spikes. Symptomatic leaf segments (from seedlings) and spikelets (from fully mature plants) were excised and surface-sterilized using 2% bleach followed by rinse with water three times. The tissues were plated onto water agar (12 g/L) supplemented with streptomycin (100 mg/L), ampicillin (100 mg/L), and kanamycin (150 mg/L) to inhibit bacterial growth. Plates were incubated at room temperature (∼22°C) under continuous light for 48 to 72 hours. Emerging colonies were incubated as described previously to induce spore production. The identity of the recovered isolates was confirmed through morphological characterization (colony appearance and spore measurements after 7-10 days) and molecular identification following Hafez et al. (2026). This was followed by repeated inoculation to confirm association with symptoms.

**Table 1:**
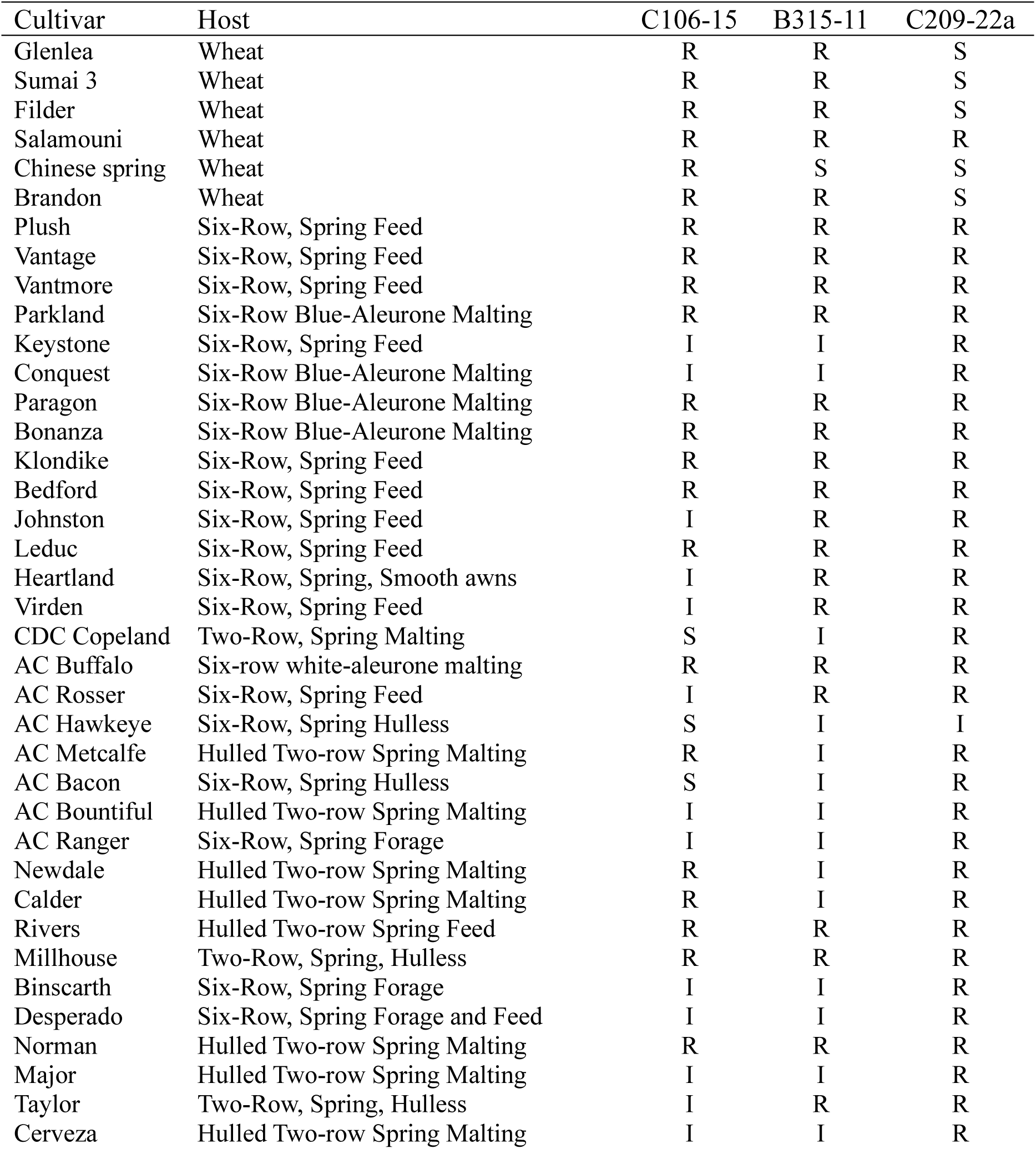
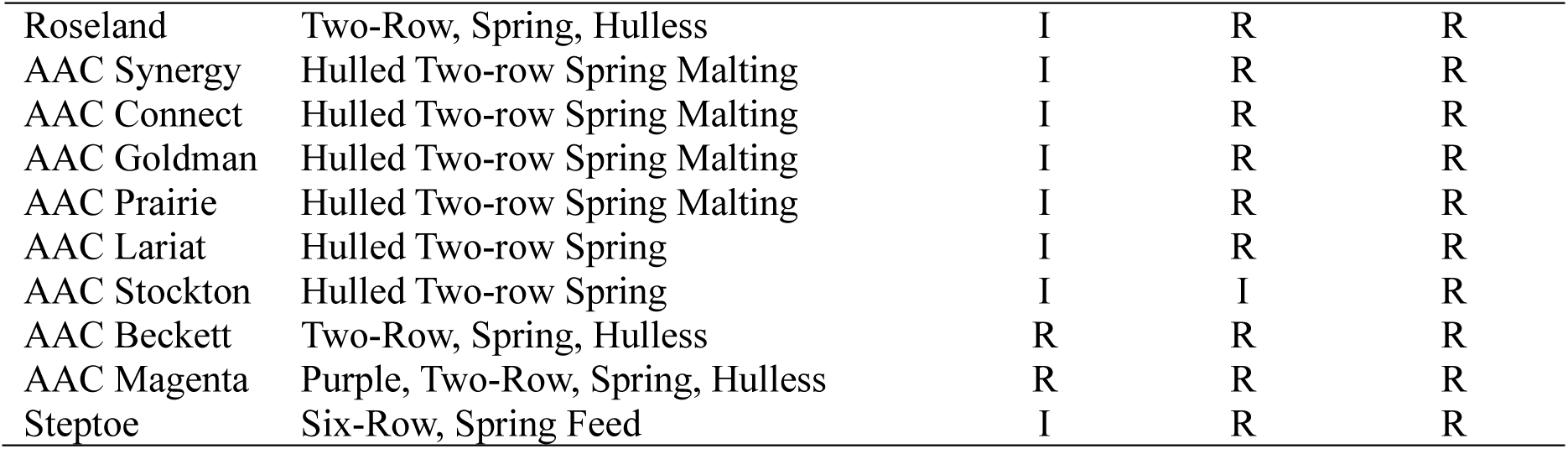
Disease reaction on leaves of wheat and barley cultivars inoculated with *P. pseudonodorum* and *P. nodorum*. Assessments were performed on the second leaf using a spore suspension of 1×10^6^ spores/mL. Wheat and barley cultivars were inoculated with *P. pseudonodorum* isolates (C106-15, B315-11) and the *P. nodorum* isolate C209-22a. Symptoms were evaluated on a 1-5 scale and classified as follows: R = Resistant (score 1-2); I = Intermediate (score 3); S = Sensitive (scores 4-5). Negative controls inoculated with sterile distilled water showed no symptoms with all tested cultivars. Ratings were derived from the average of two replicates.

| Cultivar | Host | C106-15 | B315-11 | C209-22a |
| --- | --- | --- | --- | --- |
| Glenlea | Wheat | R | R | S |
| Sumai 3 | Wheat | R | R | S |
| Filder | Wheat | R | R | S |
| Salamouni | Wheat | R | R | R |
| Chinese spring | Wheat | R | S | S |
| Brandon | Wheat | R | R | S |
| Plush | Six-Row, Spring Feed | R | R | R |
| Vantage | Six-Row, Spring Feed | R | R | R |
| Vantmore | Six-Row, Spring Feed | R | R | R |
| Parkland | Six-Row Blue-Aleurone Malting | R | R | R |
| Keystone | Six-Row, Spring Feed | I | I | R |
| Conquest | Six-Row Blue-Aleurone Malting | I | I | R |
| Paragon | Six-Row Blue-Aleurone Malting | R | R | R |
| Bonanza | Six-Row Blue-Aleurone Malting | R | R | R |
| Klondike | Six-Row, Spring Feed | R | R | R |
| Bedford | Six-Row, Spring Feed | R | R | R |
| Johnston | Six-Row, Spring Feed | I | R | R |
| Leduc | Six-Row, Spring Feed | R | R | R |
| Heartland | Six-Row, Spring, Smooth awns | I | R | R |
| Virden | Six-Row, Spring Feed | I | R | R |
| CDC Copeland | Two-Row, Spring Malting | S | I | R |
| AC Buffalo | Six-row white-aleurone malting | R | R | R |
| AC Rosser | Six-Row, Spring Feed | I | R | R |
| AC Hawkeye | Six-Row, Spring Hullless | S | I | I |
| AC Metcalfe | Hulled Two-row Spring Malting | R | I | R |
| AC Bacon | Six-Row, Spring Hullless | S | I | R |
| AC Bountiful | Hulled Two-row Spring Malting | I | I | R |
| AC Ranger | Six-Row, Spring Forage | I | I | R |
| Newdale | Hulled Two-row Spring Malting | R | I | R |
| Calder | Hulled Two-row Spring Malting | R | I | R |
| Rivers | Hulled Two-row Spring Feed | R | R | R |
| Millhouse | Two-Row, Spring, Hullless | R | R | R |
| Binscarth | Six-Row, Spring Forage | I | I | R |
| Desperado | Six-Row, Spring Forage and Feed | I | I | R |
| Norman | Hulled Two-row Spring Malting | R | R | R |
| Major | Hulled Two-row Spring Malting | I | I | R |
| Taylor | Two-Row, Spring, Hullless | I | R | R |
| Cerveza | Hulled Two-row Spring Malting | I | I | R |
| Roseland | Two-Row, Spring, Hulless | I | R | R |
| AAC Synergy | Hulled Two-row Spring Malting | I | R | R |
| AAC Connect | Hulled Two-row Spring Malting | I | R | R |
| AAC Goldman | Hulled Two-row Spring Malting | I | R | R |
| AAC Prairie | Hulled Two-row Spring Malting | I | R | R |
| AAC Lariat | Hulled Two-row Spring | I | R | R |
| AAC Stockton | Hulled Two-row Spring | I | I | R |
| AAC Beckett | Two-Row, Spring, Hulless | R | R | R |
| AAC Magenta | Purple, Two-Row, Spring, Hulless | R | R | R |
| Stephoe | Six-Row, Spring Feed | I | R | R |

### Molecular Identification of *P. pseudonodorum* and *P. nodorum*

*P. nodorum* and *P. pseudonodorum* as species identity was confirmed at two levels: *in planta* using total DNA from infected ‘Chinese Spring’ tissue, and *in vitro* using gDNA from isolates recovered from infected tissues. Identification was initially conducted by PCR using species-specific primers: Pno-Fm for *P. nodorum* and Pps-Fm for *P. pseudonodorum,* each combined with Bt-Rm universal primer, as described in Hafez et al. (2026), to specifically detect and amplify the target species. Species identities were further confirmed using PCR-RFLP assay involving amplification of a *β-tubulin* fragment using primers Bt-Fu and Bt-Ru, followed by restriction digestion using *Aat*II and *Oli*I, which differentiated between *P. nodorum* and *P. pseudonodorum*, respectively (Hafez et al. 2026).

The DNA extracted from tested fungal colonies and plant infected tissues (leaves and spikes) were used as templates in PCRs to detect *ToxA*, *SnTox1* and *SnTox3* genes to confirm there was no cross-contamination among isolates, as they display different NE profile. The primers used for these PCR are listed in Table 2. The *ToxA* amplification cycling conditions were as follows: an initial denaturation at 95°C for 2 min; followed by 30 cycles of 94°C for 50 sec, 55°C for 50 sec, 70°C for 50 sec, and a final extension at 70°C for 7 min. Cycling conditions for *SnTox1* and *SnTox3* followed the protocol described by (Gao et al. 2015) with the annealing temperature adjusted to 51°C for *SnTox1* and 60°C for *SnTox3*. Cycling conditions consisted of an initial denaturation (94°C for 3 min), followed by 35 cycles (94°C for 1 min, annealing for 50 sec, 72°C for 1 min), and a final extension (72°C for 5 min).

**Table 2:**
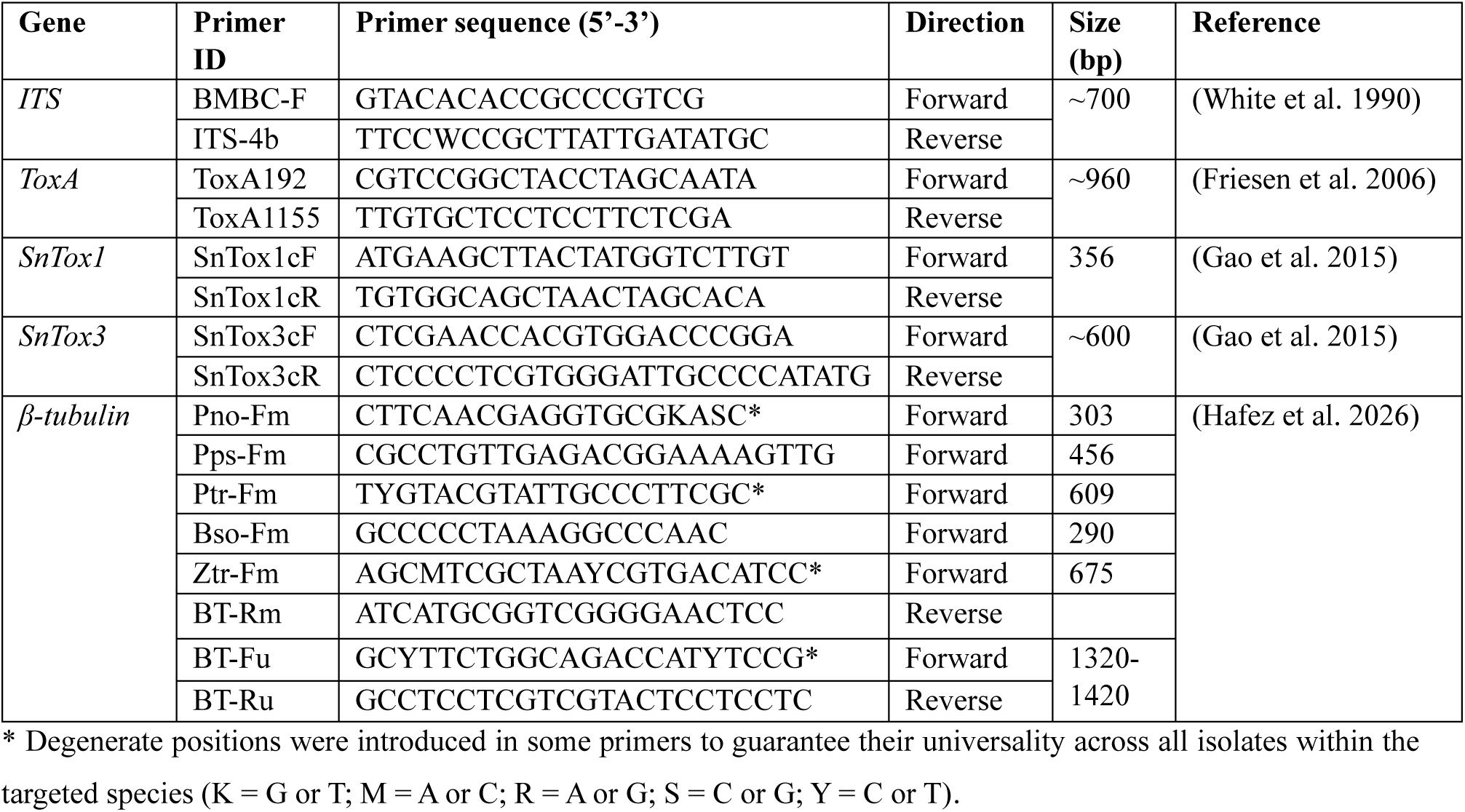
List of primers used in this study in PCR reactions.

## Results

### Symptoms development on the leaves

Symptoms development was evaluated on the second leaf of wheat and barley seedlings at 10 dpi, using a 1-5 scale (Figure 1). On wheat, the tested cultivars generally exhibited resistant reaction (R) to the *P. pseudonodorum* isolates, with the notable exception of ‘Chinese Spring’, which displayed a susceptible (S) reaction to B315-11 (Figure 2A). Conversely, the *P. nodorum* isolate C209-22a was highly virulent on most tested wheat lines, inducing reactions ranging from 3 to 5 based on the 1 to 5 scale, with 3 rated as intermediate (I), and 4 or 5 were rated as susceptible (S) on ‘Glenlea’, ‘Sumai3’, ‘Fielder’, ‘Brandon’, and ‘Chinese Spring’. Only ‘Salamouni’ was the wheat genotype rated resistant to all tested isolates. On barley, the infection pattern was reversed; all cultivars, with exception to ‘AC Hawkeye’, were rated resistant to C209-22a (*P. nodorum*), and the majority of barley cultivars were rated R or I the *P. pseudonodorum* isolates. Only three cultivars were rated S in response to C106-15. (Table 1; Figure 2B).

**Figure 1.**
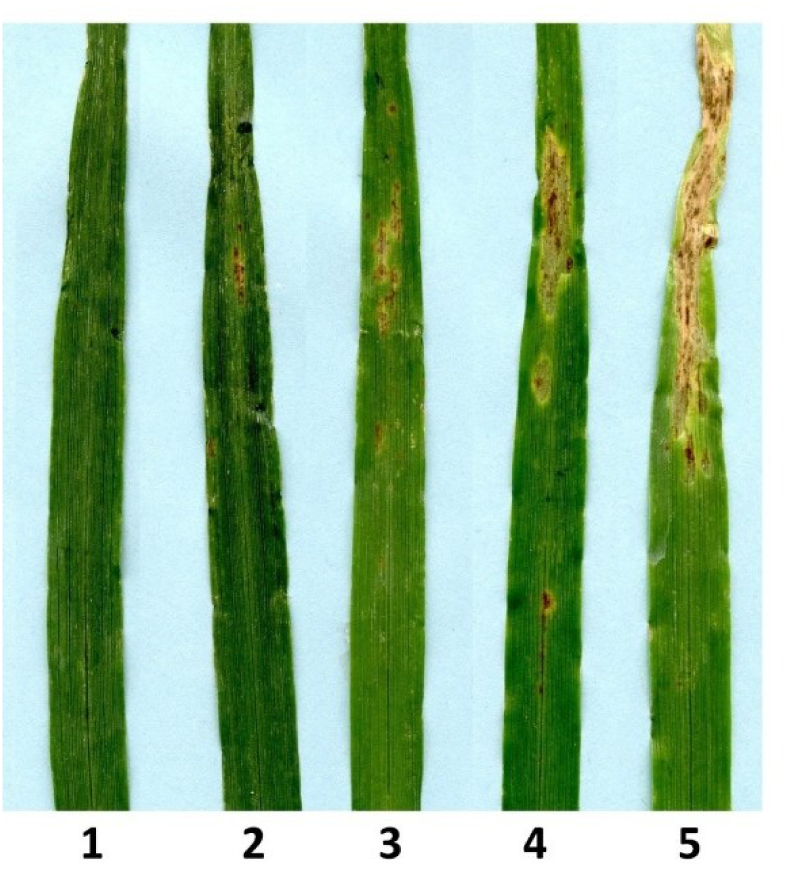
Rating scale of barley seedling response to fungal infection on the second leaf, assessed 10 days after inoculation. From left to right, leaves represent disease scores 1-5: 1,no lesion or small brown to black spots without surrounding chlorosis (resistant); 2, small brown to black spots with little chlorosis (moderately resistant); 3, small (dark) brown to black spots surrounded by a chlorotic halo (Intermediate); 4, small dark brown or black spots completely surrounded with chlorotic zones with some coalescent lesions (Susceptible); 5, dark brown or black centres could or could not be distinguishable with most lesions coalescing in chlorotic or necrotic spots (Susceptible).

**Figure 2:**
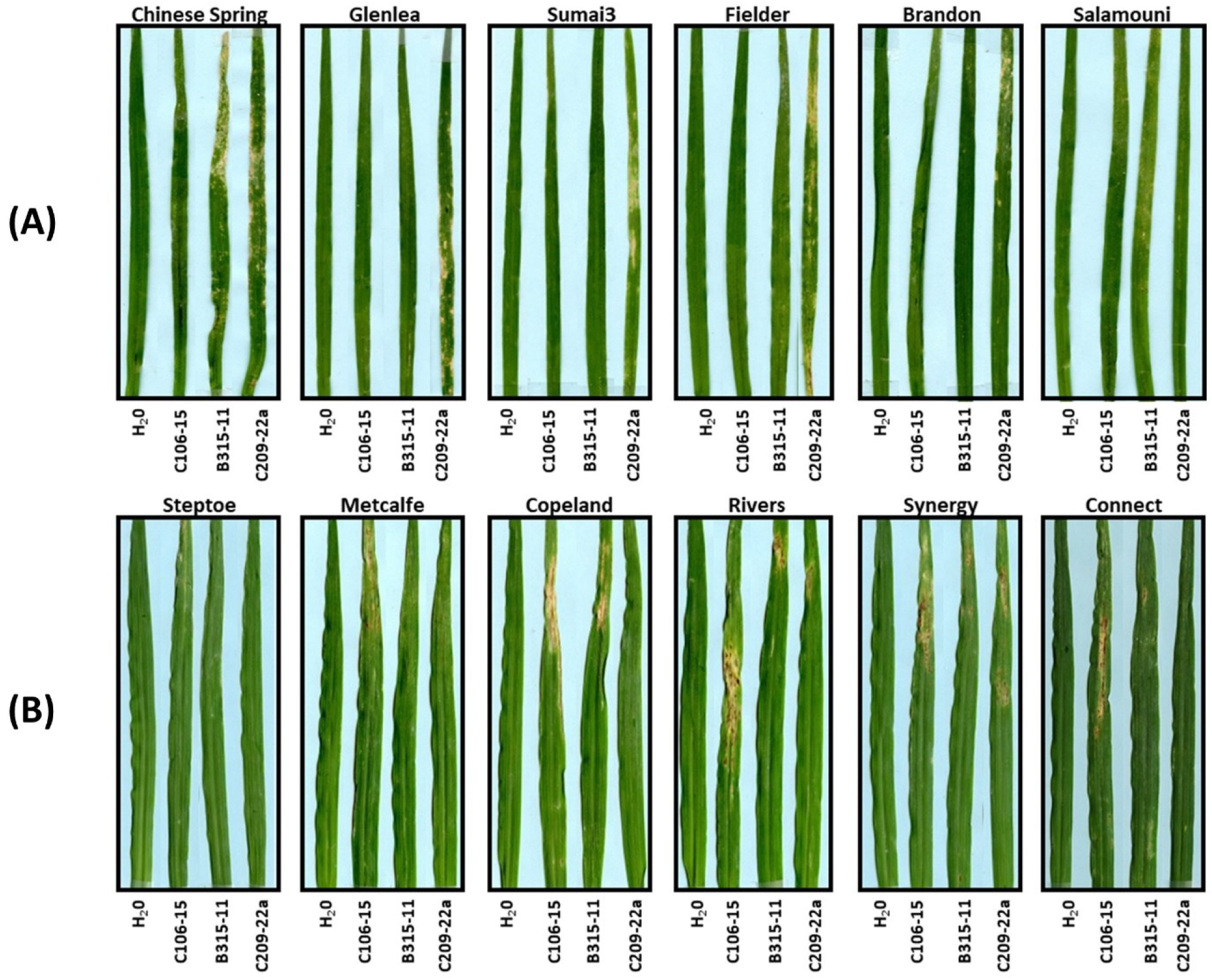
Second leaves of wheat (A) and barley (B) cultivars 10 days post-inoculation (dpi) with *P. pseudonodorum* isolates C106-15 and B315-11, or *P. nodorum* isolate C209-22a. Sterile distilled water (H_2_O) was used as the negative control. The *Parastagonospora* isolates used for the inoculation differed in necrotrophic effector profile*s*: C106-15 (no known effector), B315-11 (Tox1), and C209-22a (ToxA+, Tox1+, Tox3+).

### Symptoms development on the spikes

The ability to infect head tissues was assessed by the percentage of infected spikelets in each spike (Table 3), and by the severity of developed symptoms, evaluated by the intensity of necrotic lesions, on a subset of selected cultivars of both wheat and barley (Figure 3). In general, the *P. pseudonodorum* isolates (C106-15 and B315-11) induced over 40% of infection on wheat (Table 3), but symptom severity on infected spikelets were weak (Figure 3A). In the two barley genotypes, infection ranged from 21% to 82% with considerably more severe symptoms on ‘Rivers” than on ‘Steptoe’ and wheat (Table 3 Figure 3A, B).

**Figure 3:**
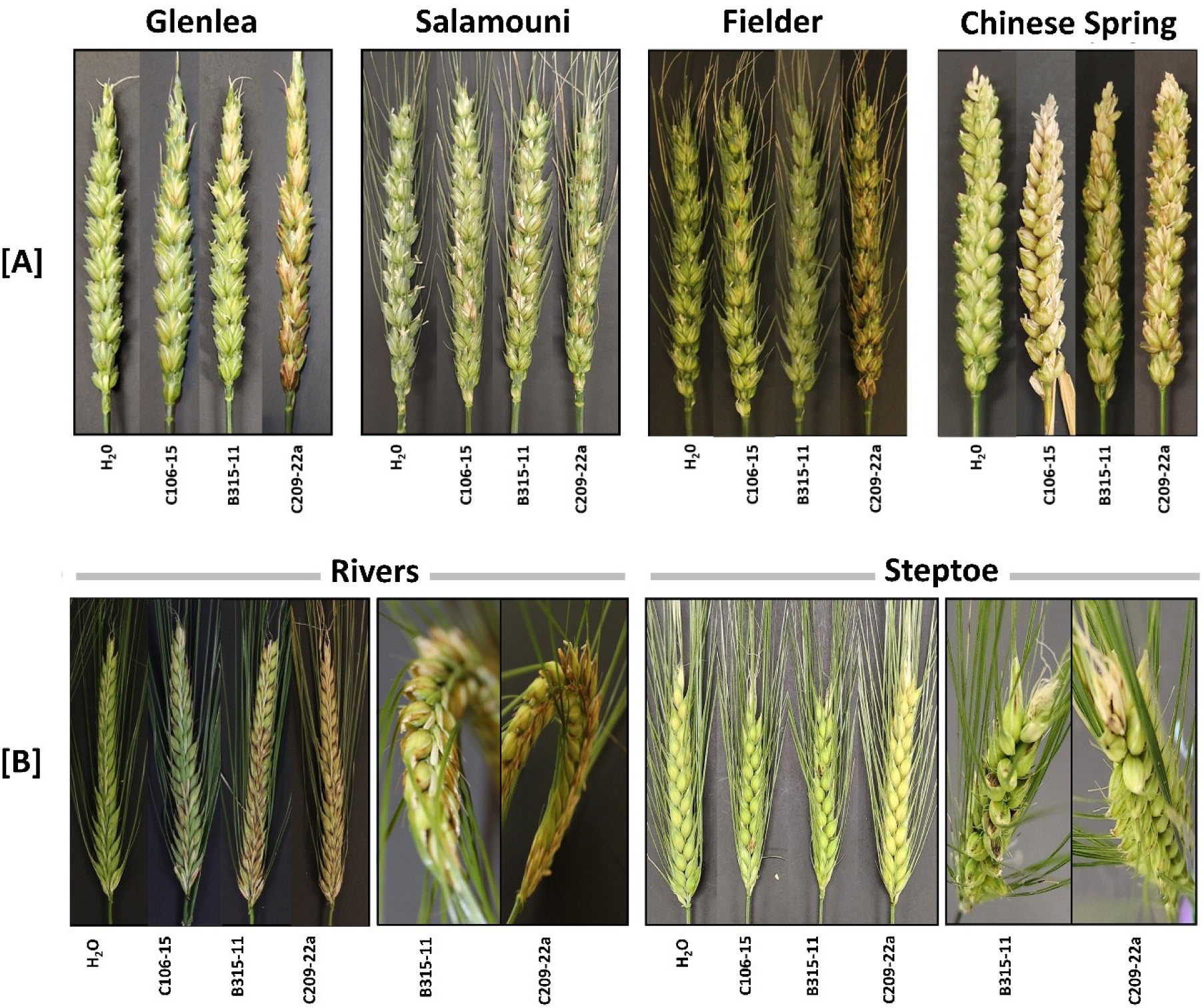
**[A] Disease symptoms on wheat spikes 18 dpi.** Spikes inoculated with *Parastagonospora* isolates: *P. pseudonodorum* (C106-15 and B315-11) or *P. nodorum* (C209-22a). Sterile distilled water (H_2_O) was used as the mock control. Symptom morphology on wheat spikelets varies by pathogen species*: P. pseudonodorum* infection resulted in brownish lesions with lighter margins, whereas *P. nodorum* typically produced more extensive, dark, and purplish necrotic lesions. **[B] Symptom development on barley spikes 18 dpi.** Spikes were inoculated with *P. pseudonodorum* (C106-15, B315-11) or *P. nodorum* (C209-22a), with sterile distilled water (H₂O) used as a mock control. Although the closed architecture of the barley spike initially obscured symptom visibility, internal examination of the spikelets revealed strong, evident infection across the inoculated samples.

**Table 3:** Disease reaction on infected wheat and barley spikes inoculated with *P. pseudonodorum* and *P. nodorum*. Inoculations were performed on heads during anthesis using a spore suspension of 1×10^6^ spores/mL. Four wheat cultivars (Glenlea, Salamouni, Fielder, Chinese Spring) and two barley cultivars (Steptoe, Rivers) were inoculated with *P. pseudonodorum* (C106-15 and B315-11) and the *P. nodorum* isolate C209-22a. Ability to infect was evaluated as the percentage of infected spikelets per head. Values represent the average of two replicates. Negative controls inoculated with sterile distilled water showed no symptoms across all tested cultivars.

| Cultivar | Host | C106-15 | B315-11 | C209-22a |
| --- | --- | --- | --- | --- |
| Glenlea | Wheat | 53% | 65% | 98% |
| Salamouni | Wheat | 86% | 99% | 96% |
| Fielder | Wheat | 40% | 61% | 75% |
| Chinese Spring | Wheat | 83% | 84% | 95% |
| Steptoe | Barley | 21% | 37% | 18% |
| Rivers | Barley | 82% | 80% | 80% |

Clear isolate x genotype interaction was observed, with B315-11 generally causing higher infection rates and greater symptom severity than C106-15. In contrast, the *P. nodorum* isolate, C209-22a, caused higher infection overall on wheat than *P. pseudonodorum*, whereas infection percentages on barley were similar to those induced by *P. pseudonodorum*. Symptom severity on ‘Steptoe’ was higher following infection with *P. pseudonodorum* than with *P. nodorum* (Figure 3B). Although the Steptoe spikes appeared largely asymptomatic when viewed without manipulation, the closed structure of the barley spikelets obscured the visualization of infection. Bending the spike or gently separating the spikelets to expose their basal portions revealed distinct dark brown to black lesions on the glumes extending to the rachis (Figure 3B). Symptoms following *P. pseudonodorum* infection became visible on the heads between 4 and 8 dpi, and were frequently accompanied by chlorotic or necrotic lesions on the stems below the infected spikes, particularly in ‘Rivers’ inoculated with B315-11. By 18 dpi, no considerable differences in the percentage of infected spikelets were observed among ‘Salamouni’, ‘Fielder’, or ‘Chinese Spring’ across the three fungal isolates (Figure 4). However, on ‘Glenlea’, *P. nodorum* (C209-22a) was significantly more aggressive, causing 98% infection, nearly double the infection rates (53-65%) observed with the *P. pseudonodorum* isolates (Figure 3A Table 3).

**Figure 4:**
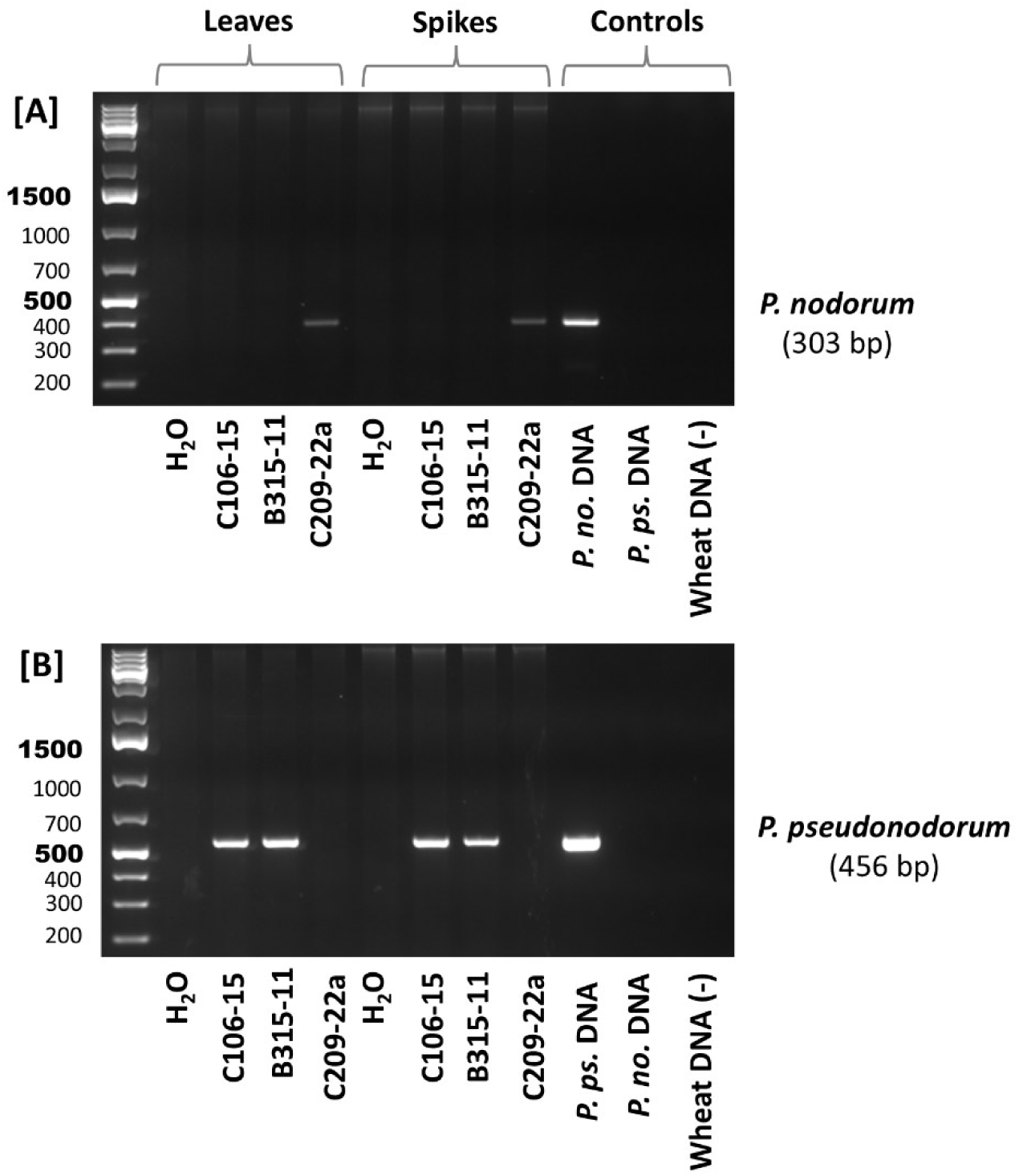
Specific detection of *Parastagonospora* species in inoculated wheat tissues. DNA extracted from “Chinese Spring” wheat seedling leaves and spikes served as the template for PCR amplification of the *β-tubulin* (*tub1*) gene region using species-specific primer sets. [A] Specific detection of *P. nodorum* (303 bp) in tissues inoculated with isolate C209-22a. [B] Specific detection of *P. pseudonodorum* (456 bp) in tissues inoculated with isolates C106-15 and B315-11. Controls include genomic DNA from pure cultures (*P. no.* DNA and *P. ps.* DNA), non-inoculated wheat DNA, and negative control for PCR (NC).

### Molecular detection and confirmation of pathogenicity on leaves and spikes

*In planta* validation using infected ‘Chinese Spring’ leaves and spikelets confirmed the causal agents and its association with the developed symptoms. The presence of *P. nodorum* and *P. pseudonodorum* was conclusively verified through molecular characterization. To assess the colonization and presence of *Parastagonospora* species within host tissues, we performed PCR-based diagnostic assays targeting the *β-tubulin* (*tub1*) gene region using genomic DNA extracted from the leaves and spikes of ‘Chinese Spring’ wheat (Figure 4). This assay confirmed the species-specific detection of *P. nodorum* and *P. pseudonodorum* through the generation of distinct amplicons: leaf and spike tissues inoculated with *P. nodorum* isolate C209-22a yielded a 303 bp fragment using Pno-Fm/Bt-Rm specific primers (Figure 4A), whereas no amplicons obtained in tissues inoculated with *P. pseudonodorum* isolates (C106-15 and B315-11). On the other hand, tissues inoculated with C106-15 and B315-11 produced a 456 bp fragment using Pps-Fm/Bt-Rm specific primers (Figure 4B), whereas no amplicons obtained in tissues inoculated with C209-22a. These results were validated by comparing them against positive controls derived from the DNA of single-spore cultures of the respective pathogens, *P. nodorum* and *P. pseudonodorum*, and negative controls consisting of non-inoculated wheat DNA. The absence of amplification in the non-inoculated samples, combined with the precise matching of amplicon sizes to the pure culture controls, confirms the specificity and reliability of the *tub1*-based primer set for identifying and differentiating *P. nodorum* and *P. pseudonodorum* in colonized wheat tissue.

In addition, PCR-RFLP assays specific for *P. nodorum* and *P. pseudonodorum* were conducted on DNA extracted from ‘Chinese Spring’ infected leaf and spike tissues. The universal *tub1* gene specific primers, Bt-Fu and Bt-Ru, amplified a 1320 bp fragment from the leaf and spike tissues inoculated with C106-15, B315-11, and C209-22a (Figure 5A). Digesting the amplified PCR fragments with *Aat*II (specific to *P. nodorum*) cleave PCR fragments in leaf and spike tissue inoculated with C209-22a. The restriction digestion produces a pattern specific to *P. nodorum* that consist of 354 bp and 966 bp; however, no cleavage on tissues inoculated with C106-15 and B315-11 were observed (Figure 5B). Similarly, digesting the amplified PCR fragments with *Oli*I (specific to *P. pseudonodorum*) cleaves PCR fragments in leaf and spike tissue inoculated with C106-15 and B315-11 isolates. The restriction digestion generates a pattern specific to *P. pseudonodorum* that consist of 533 bp and 787 bp, with no cleavage on tissues inoculated with the C209-22a *P. nodorum* isolate (Figure 5C).

**Figure 5.**
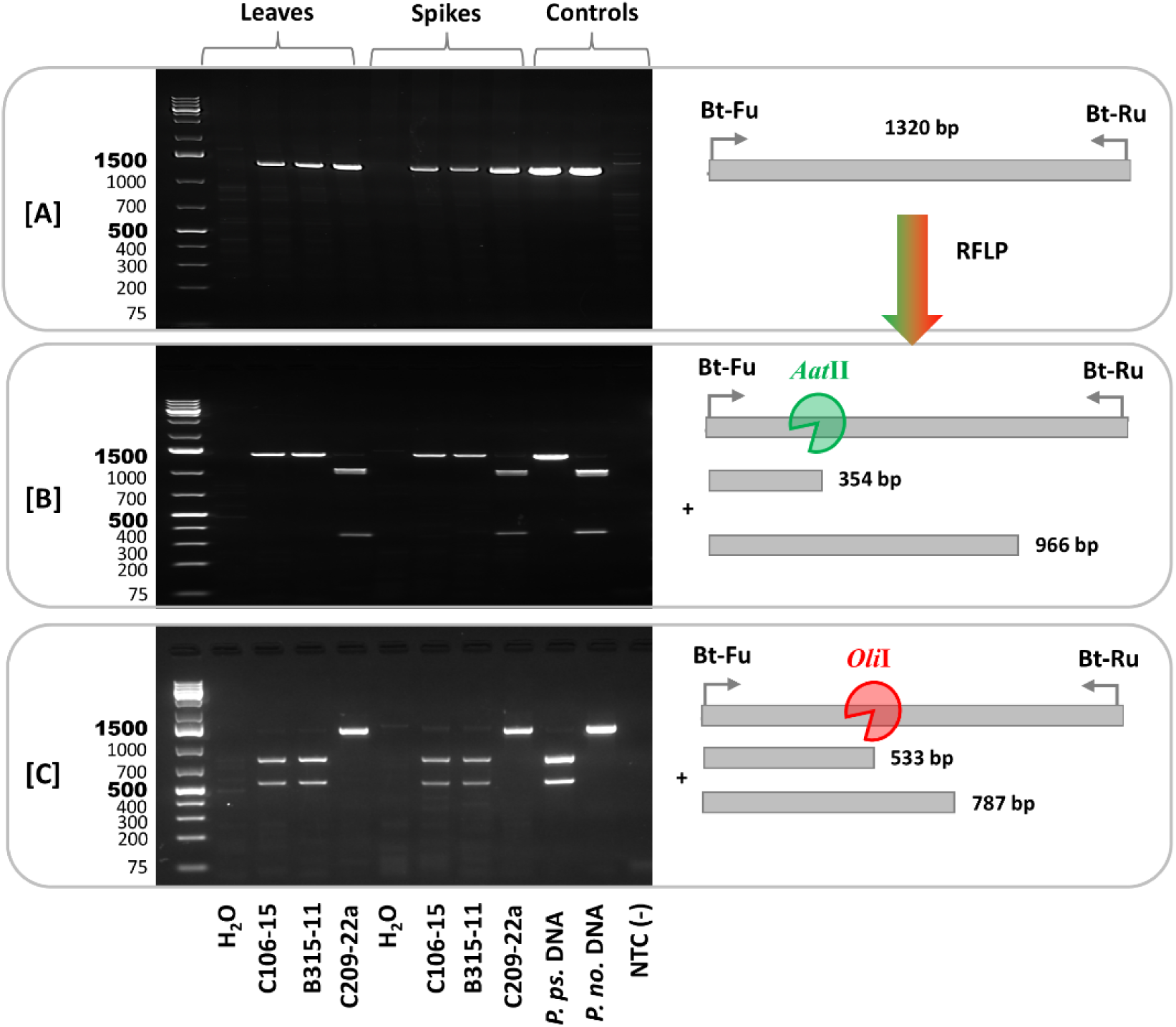
Identification of *Parastagonospora* species in inoculated Chinese Spring wheat tissues via PCR-RFLP. **[A]** Amplification of a 1320 bp region of the β-tubulin (*tub1*) gene from infected leaf and spike tissues using primers Bt-Fu and Bt-Ru. **[B]** Restriction digestion of the PCR product with the *Aat*II enzyme, which specifically cleaves *P. nodorum* DNA into 354 bp and 966 bp fragments. **[C]** Restriction digestion with the *Oli*I enzyme, which specifically cleaves *P. pseudonodorum* DNA into 533 bp and 787 bp fragments. Controls include genomic DNA from pure cultures (*P. no.* DNA and *P. ps.* DNA), non-inoculated wheat DNA, and negative control for PCR (NC).

Re-isolation of the pathogen from symptomatic tissues consistently yielded the same isolates used in the inoculation, and re-inoculation reproduced the expected disease symptoms. The recovered isolates showed distinct colony morphologies that matched their original identities (Figure 6). *P. pseudonodorum* isolate C106-15 produced a cream-coloured colony with a darker gray center, whereas isolate B315-11 formed a pink to orange colony. The *P. nodorum* isolate C209-22a produced a pinkish colony that became brownish over time and occasionally developed a dark halo (Figure 6A). Spore morphology and measurements were also consistent with the expected features of each isolate. ITS sequences amplified from the re-isolated pathogens (∼700 bp; Figure 6B) from spike tissue of each treatment showed identical sequence to the original species used for inducing the infection. In addition, PCR assays of the NE genes matched the original effector profiles of the inoculated strains: C106-15 (no effector), B315-11 (*Tox1*), and C209-22a (*ToxA*, *Tox1*, *Tox3*). In particular, *ToxA* (960 bp) was detected in the mycelia re-isolated from the spikes of ‘Rivers’ and ‘Salamouni’ inoculated with the *P. nodorum* C209-22a isolate (Figure 6C). *Tox1* (350 bp) was detected in the mycelia obtained from ‘Rivers’, ‘Glenlea’, and ‘Salamouni’ spikes inoculated with the *P. nodorum* C209-22a and the *P. pseudonodorum* B315-11 isolates (Figure 6D). *Tox3* (600 bp) was detected in the mycelia from ‘Rivers’, ‘Glenlea’, and ‘Salamouni’ spikes inoculated with the C209-22a isolate (Figure 6E). Although these results only showed three cereal cultivars, it is only a representation of all samples tested, and all exactly matched the NE profile of the inoculated pathogen. Together, these results confirmed that the pathogens recovered from infected tissues were the same as those originally used for inoculation.

**Figure 6.**
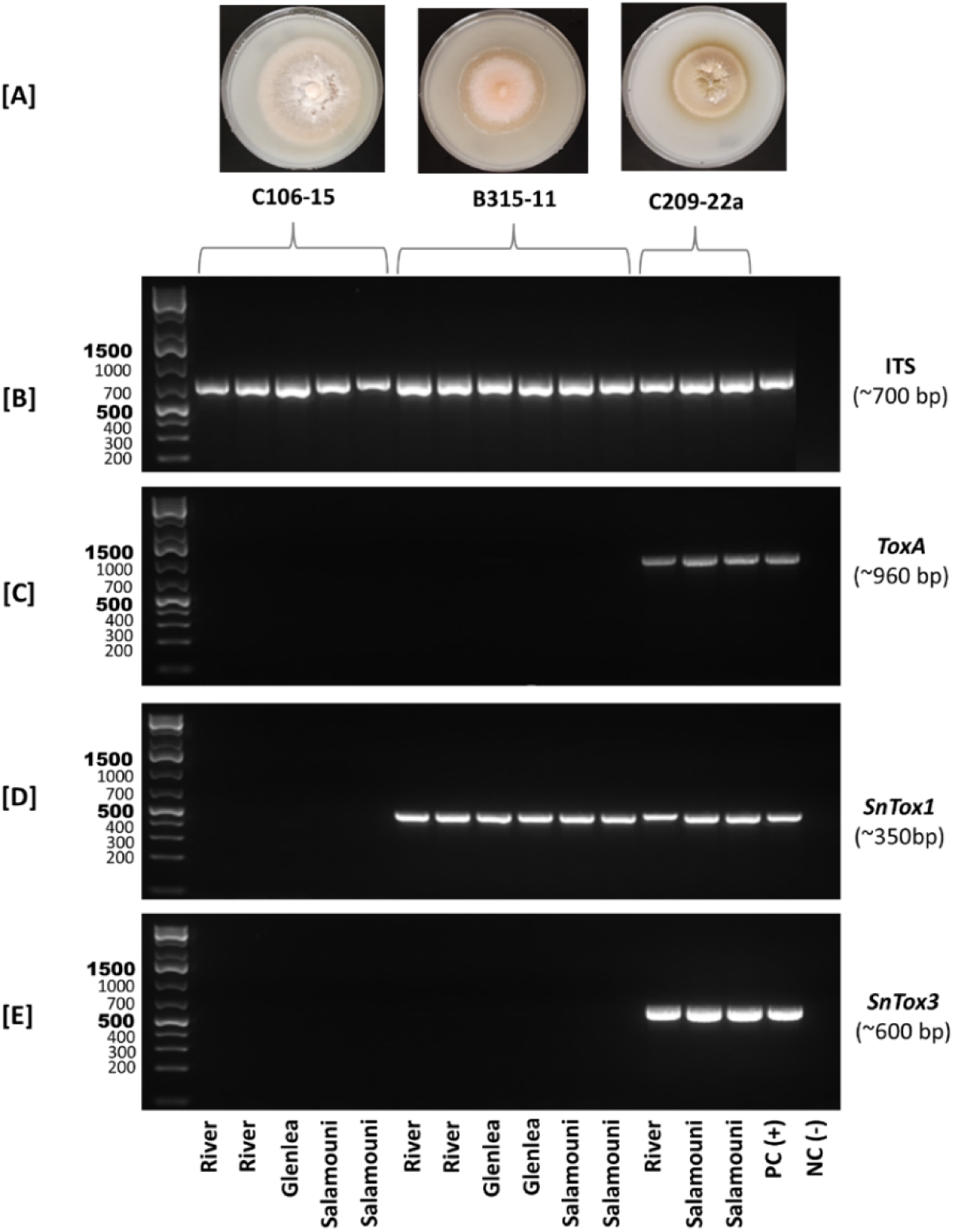
Molecular confirmation of *Parastagonospora* species re-isolated from inoculated wheat (Glenlea and Salamouni) and barley (Rivers) spikes to satisfy Koch’s postulates. [A] Morphological characterization of re-isolated fungal colonies from *P. pseudonodorum* (isolates C106-15 and B315-11) and *P. nodorum* (isolate C209-22a), showing colony phenotypes consistent with the original inocula. [B-E] PCR-based molecular identification of the re-isolated colonies. Species identity was confirmed by amplification of the Internal Transcribed Spacer (ITS) region (∼700 bp) across all samples [B]. effector gene profiling confirmed the distinct identities of the isolates: [C] *SnToxA* (∼960 bp) was detected exclusively in *P. nodorum* C209-22a; [D] *SnTox1* (∼350 bp) was detected in all *P. pseudonodorum* B315-11 and *P. nodorum* C209-22a samples; and [E] *SnTox3* (∼600 bp) was detected specifically in the *P. nodorum* C209-22a isolate. Positive control, PC (+), utilized genomic DNA from a pure culture of *P. nodorum* C209-22a, while the negative control, NC (-) of the PCR reaction was distilled water. Host cultivars (Rivers, Glenlea, and Salamouni) are indicated below the gel images.

Collectively, these molecular results, in combination with the fulfillment of Koch’s postulates, confirm that the observed symptoms were directly attributable to the inoculated pathogen.

## Discussion

In this study, we addressed a gap in our understanding of *P. pseudonodorum* pathogenicity on wheat and barley, confirming its ability to infect the leaves and spikes of both hosts. However, symptoms were generally weak on the leaves of both hosts and on wheat spikes, whereas more evident damage was observed on barley spikes. However, we tested only a limited number of isolates and host genotypes as a proof of concept; therefore, future studies with a broader range of isolates and genotypes will be needed. This study represents the first modern validation of *P. pseudonodorum* pathogenicity using two well-characterized isolates, one of which has a fully assembled genome, providing much needed clarity regarding the long-standing ambiguity surrounding this species and its pathogenicity.

Following the first identification of *P. pseudonodorum* (Pat1) in 1942 (Johnson, 1947), the literature has reported contradictory findings regarding whether this species causes greater damage to wheat or barley, whether it occurs more frequently on leaves or grains, and whether it is more prevalent on wheat than on barley (Cunfer 2000; Johnson 1947; Richardson and Noble, 1970; Shaw 1957a; Shipton et al. 1971). Although subsequent studies on its genetic diversity, genomic characteristic, and phylogenetic relationship and related lineages (Croll et al. 2023; Ghaderi et al. 2022; McDonald et al. 2012; Ueng et al. 1994; Ueng et al. 1998; Ueng et al. 2003; Zhang et al. 2009), we remain unaware of recent studies that systematically reassessed its pathogenicity through controlled inoculation experiments using molecularly confirmed isolates. Earlier reports indicated that *P. pseudonodorum* was more common in Western Canada, whereas *P. nodorum* was more common in Eastern Canada (Johnson 1947). Its prevalence has also been reported to fluctuate from trace to widespread, depending on the year and the host (Cunfer 2000; Johnson 1947; King et al. 1983; Shipton et al. 1971). These inconsistencies are not surprising given the inconspicuous nature of the symptoms, the similarity in spore morphology among these species, and the lack of specific molecular diagnostic tools. While some early reports described *P. pseudonodorum* as a weak pathogen of wheat, but they noted its ability to cause damage later in the season following prolonged latent period (Johnson 1947; Shaw 1957a). Other studies reported that it was unable to infect grains, whereas others reached the opposite conclusion (Shipton et al. 1971).

In this study, inoculation of 42 barley genotypes with *P. pseudonodorum* showed that most of lines exhibited resistance to intermediated reactions following leaf infection, indicating limited disease ability on the leaves. On the other hand, inoculation on a limited number of wheat and barley genotypes at the anthesis stage, showed a higher level of symptom development that varied depending on either the isolate or the cultivar. The symptoms severity were not necessarily associated with the effectors produced by a certain isolate, as ‘Salamouni’ which tested resistant to all isolates at the seedling stage exhibited a higher % of infection on the spikes when inoculated with these isolates. Previous studies have suggested a possible association between *P. pseudonodorum* and tissues specificity based on its reported association with grains rather than leaves (Hafez et al. 2020; McDonald et al. 2012). However, such an assumption need a revisit, as by default these studies relied on isolates that were not obtained through a random survey designed to represent grain and leaf samples equally, and therefore the reported association with grains rather than leaves does not conclusively establish tissue specificity.

Despite the worldwide prevalence of *P. pseudonodorum* on cereals and its potential impact on glum blotch, there is a limited interest from the research community, most likely due to the limited availability of developed diagnostic tools to accurately differentiate it from *P. nodorum* (Hafez et al. 2026). This is reflected in the 50-fold higher number of research hits for *P. nodorum* compared to *P. pseudonodorum* (Pat1) (62000 *vs* 1200). However, *P. pseudonodorum* is gaining more interest in recent years given that it is considered a quarantine species in countries, such as China (Zhao 2021). The ability of *P. pseudonodorum* to infect barley have been observed in Iran (Ghaderi et al. 2022), but there is no published experimental validation for those statements. *P. pseudonodorum* was first characterized in Canada and was considered a forma speciales of *Leptosphaeria avenaria* and was named *Leptosphaeria avenaria* f. sp. *triticea* (Johnson 1947). It was then reported as a wheat pathogen, and a weak pathogen of barley and rye (Johnson 1947), but later more reports on it from the UK linked it to barley, including an epidemic in the 1970’s in Scotland (Richardson and Noble, 1970). In this current study, despite the limited number of wheat and barley genotypes screened, we have found strong proof of its impact on spike tissue rather than on the leaves. However, further work is needed to evaluate this on a wider selection of genotypes and isolates.

In terms of prevalence and occurrence, many samples collected in the 1940s had both *P. nodorum* and *P. pseudonodorum*. Although the two could not be distinguished based solely on visual inspection of symptoms on infected leaves, but scraping spores from infected leaves showed the presence of both species on same infected leaf (Johnson, 1947). In recent works the Pat1 isolates were obtained mainly from grain samples (Hafez et al. 2020; McDonald et al. 2012), in contrast to *P. nodorum* which predominates the node samples (Hafez et al. 2020). However, a recent survey in Western Canada showed that *P. pseudonodorum* is widespread on wheat leaves as well (unpublished data).

Although *P. pseudonodorum* was first identified in Canada in 1942, under an earlier taxonomic name (Johnson 1947), nevertheless, it was likely not a newly emerged species. Examination of herbarium specimens collected in Manitoba (MB) and Saskatchewan (SK) in 1927 and 1928 suggested its presence, supported by notes indicating spore sizes that differed from those typically observed for *Septoria* (reported in (Johnson 1947). However, *P. pseudonodorum* was not officially reported as a new *forma specialis* of *Phaeosphaeria avenaria* until 1947. Between 1943 to 1946 it was found in Western Canada (MB, SK and Alberta (AB)) and Eastern Canada (Ontario (ON), Quebec (QC), and the Maritime provinces) on wheat, barley, oat and other grasses in the *Elymus* and *Agropyron* species (Johnson 1947).

In conclusion, while *P. pseudonodorum* is of a worldwide distribution across wheat growing regions, its taxonomic ambiguities have historically hindered accurate research into its biological role. By addressing key gaps in its diagnosis and pathogenicity across leaves and spiked using well characterized isolates, this study clarifies long standing uncertainties. Given its critical role in head infection and its status as a quarantine pathogen of phytosanitary concern, these findings are essential for future epidemiological surveillance, seed screening and host-pathogen interaction studies.

## Acknowledgments

Thanks to Dr. Ana Badea for providing us with the barley cultivars and the valuable discussion on this work, and thanks to Dr. Kelly Turkington for providing much scientific discussion on septoria and barley and some of the barley genotypes as well. Thanks for Dr. Timothy Friesen for providing protocols on Parastagonospora and some confirmation to the effectors genes.

## Fund

This work was funded by various projects from AB Grains, Saskatchewan Wheat Development SWD, Result Driven Agriculture Research (RDAR), and Agriculture and Agri-Food Canada to Dr. Aboukhaddour’s Lab

## Notes

### Competing Interest Statement

The authors have declared no competing interest.

